# Oligomerisation of a plant helper NLR requires cell-surface and intracellular immune receptor activation

**DOI:** 10.1101/2022.06.16.496440

**Authors:** Joanna M. Feehan, Junli Wang, Xinhua Sun, Jihyeon Choi, Hee-Kyung Ahn, Bruno Pok Man Ngou, Jane E. Parker, Jonathan D. G. Jones

## Abstract

Plant disease resistance involves both detection of microbial molecular patterns by cell-surface pattern recognition receptors and detection of pathogen effectors by intracellular NLR immune receptors. NLRs are classified as sensor NLRs, involved in effector detection, or helper NLRs required for sensor NLR signalling. TIR-domain-containing sensor NLRs (TNLs) require helper NLRs NRG1 and ADR1 for resistance, and their activation of defense also requires the lipase-domain proteins EDS1, SAG101 and PAD4. We investigated how the helper NLR NRG1 supports TNL-initiated immunity with EDS1 and SAG101. We find that NRG1 associates with EDS1 and SAG101 at the plasma membrane and in the nucleus, but only self-associates at the plasma membrane. Activation of TNLs is sufficient to trigger NRG1-EDS1-SAG101 interaction, but cell-surface receptor-initiated defense is also required to form an oligomeric Resistosome. The data point to formation of NRG1-EDS1-SAG101 heterotrimers in the nucleus upon intracellular receptor activation alone and indicate formation of NRG1-EDS1-SAG101 Resistosomes at the plasma membrane upon co-activation of intracellular and cell surface-receptor pathways.

## Introduction

Plants have powerful mechanisms that thwart attempted pathogen ingress and attenuate disease, but to be effective they must be activated quickly. Rapid defense induction depends on pathogen recognition, which is achieved by both cell surface and intracellular immune receptors that detect pathogen-derived molecules and initiate signalling that activates defense mechanisms (1).

Most plant disease *Resistance* (*R*) genes encode intracellular “NLR” immune receptors that carry a nucleotide-binding domain and a leucine-rich repeat domain. Many sensor NLRs that directly or indirectly detect pathogen effector molecules also require “helper” NLR(s) for their signalling (2). Most sensor NLRs carry either a TIR (Toll/Interleukin-1 receptor/Resistance) domain or a CC (coiled-coil) domain at their N-termini. All characterized TIR-NLRs (TNLs) require the helper NLR NRG1 and/or ADR1 sub-families for their full function (3). Structures of activated TNLs (4, 5) and CC-NLRs (CNLs) (6–8) reveal formation, respectively, of tetrameric or pentameric “Resistosomes” upon immune activation. TNL Resistosomes activate signalling via a TIR domain-dependent NADase activity (5), while ZAR1 Resistosomes initiate defense signaling at the plasma membrane (PM) via cation channel formation (9).

The NRG1 and ADR1 sub-families of helper NLRs and EDS1-family lipase-like proteins work together in two distinct nodes to mediate defense (10–12). In Arabidopsis, three genetically redundant ADR1s (ADR1, ADR1-L1 and ADR1-L2), co-function with EDS1 and PAD4 in potentiating transcriptional defenses which restrict pathogen growth, while NRG1s (NRG1.1 and NRG1.2) co-function with EDS1 and SAG101 to execute a hypersensitive cell death response (HR). The NRG1 and ADR1 modules can partially substitute for the other (12, 13). In *Nicotiana benthamiana*, the NRG1-EDS1-SAG101 module functions in both cell death and transcription, while the contribution of ADR1-EDS1-PAD4 remains unclear (11). Arabidopsis EDS1-PAD4 and EDS1-SAG101 dimers are receptors that bind distinct TIR enzyme-derived small molecules which allosterically induce their direct associations, respectively, with ADR1 and NRG1 helper NLRs (14, 15). These data provide a biochemical mechanism for activation of the two EDS1 – helper NLR modules but it remains unknown how the induced complexes then function to activate transcription and cell death.

Previously, we found that Arabidopsis NRG1 associates with EDS1 and SAG101 upon effector delivery in Arabidopsis (12). When transiently over-expressed in *N. benthamiana, At*NRG1.1 localizes to endomembrane networks (16, 17) while auto-active *At*NRG1A^D485V^ becomes localized to the PM to form a calcium-permeable cation channel potentially similar to ZAR1 (16). In contrast, Arabidopsis EDS1-SAG101 dimers primarily localize to the nucleus (11, 18). Yet, there is evidence for NRG1-EDS1-SAG101 associations at nuclei, or their periphery, in immune-activated tissue (12), and a recent report found small pools of EDS1, PAD4, ADR1 and SAG101 in close proximity to PM-localized SOBIR1 receptors (19). These reports illustrate the need for clear data that define the sub-cellular compartments(s) in which NRG1, EDS1, and SAG101 co-implement their immune functions upon defense activation.

Both cell surface and intracellular immune receptor co-activation are required to mount full defense responses in plants (20, 21). Even though cell-surface and intracellular responses are mediated by distinct classes of immune receptors, they mutually potentiate shared pathways. ADR1 has been reported to play a role in cell surface receptor responses (19, 22); however, no such role was reported for NRG1.

Here we show that effector recognition in Arabidopsis is required for NRG1 self-association and formation of a high molecular weight NRG1-containing oligomer, though only a small proportion of NRG1 protomers become incorporated into this oligomer. We also show that NRG1 associates with EDS1 and SAG101 at the PM and in the nucleus, but only self-associates at the PM upon effector recognition. Although NRG1 can associate with EDS1 and SAG101 upon TNL activation alone, we find that both cell surface receptor and intracellular TNL activation are required for the formation of a high molecular weight NRG1-EDS1-SAG101 complex, consistent with Resistosome oligomers. Collectively, these data reveal the formation of a putative NRG1-EDS1-SAG101 heterotrimer in nuclei, induced by TNL activation alone, and a putative NRG1-EDS1-SAG101 Resistosome at the PM requiring cell surface-receptor co-activation with a TNL intracellular receptor.

## Results

### NRG1 associates with EDS1 and SAG101 at the plasma membrane and nucleus upon effector-dependent defense activation

We previously reported that NRG1 associates with EDS1 and SAG101 upon effector recognition in Arabidopsis (12). To investigate further the role of these components in TNL-mediated immune signalling, we generated stable transgenic lines in a Col-0 *nrg1*.*1 nrg1*.*2 sag101-3* mutant background (12) carrying complementing pNRG1.2:NRG1.2-FLAG, pSAG101:SAG101-HA, and pEDS1:EDS1-V5 (*SI Appendix*, Fig. S1a & S1b). As previously (12), we utilized the Pf0-1 EtHAn system (23) (hereafter referred to as “Pf0-1”) for delivery of effector AvrRps4 that is recognized by the endogenous Arabidopsis TNL pairs RRS1/RPS4 and RRS1B/RPS4B (24) to test induction of NRG1 association with EDS1 and SAG101.

Leaves of transgenic Arabidopsis carrying epitope-tagged NRG1, EDS1 and SAG101 were infiltrated with Pf0-1 carrying either AvrRps4 or non-recognized AvrRps4^EEAA^ (25) or were mock-infiltrated with MgCl_2_ before harvest 8 hours post-infiltration (hpi) and immunoprecipitation (IP). Non-infiltrated leaves were harvested as pre-activation control. We observed a previously unnoticed weak, constitutive EDS1-V5 and SAG101-HA association with NRG1.2-FLAG (Fig. 1a). We infer that the weak association in these Arabidopsis lines is due to a higher sensitivity of α-HA and α-V5 epitope tags compared to earlier studies. Pf0-1 AvrRps4^EEAA^- and MgCl_2_-infiltrated leaves showed a slight enhancement of EDS1-V5 and SAG101-HA association with NRG1.2-FLAG compared to un-infiltrated control (Fig. 1a). These data show that cell-surface receptor activation weakly induces EDS1 and SAG101 association with NRG1 (Fig. 1a). While this was not previously observed for Arabidopsis assays, transient *N. benthamiana* Agro-infiltration assays, which likely activate cell surface receptors, trigger a weak NRG1 association with EDS1 and SAG101 (12). We verified our previous finding (12) that EDS1-V5 and SAG101-HA association with NRG1.2-FLAG is strongly enhanced upon effector delivery (Fig. 1a). The early NRG1 interaction with EDS1 and SAG101 (4 hpi), which accumulates over time as observed in transient *N. benthamiana assays* (12), is also detected in stable Arabidopsis lines (Fig. 1b). Collectively, these data suggest there may be a dynamic on-off equilibrium of NRG1 with EDS1 and SAG101 that is stabilized by TNL-mediated effector recognition.

**Figure 1.**
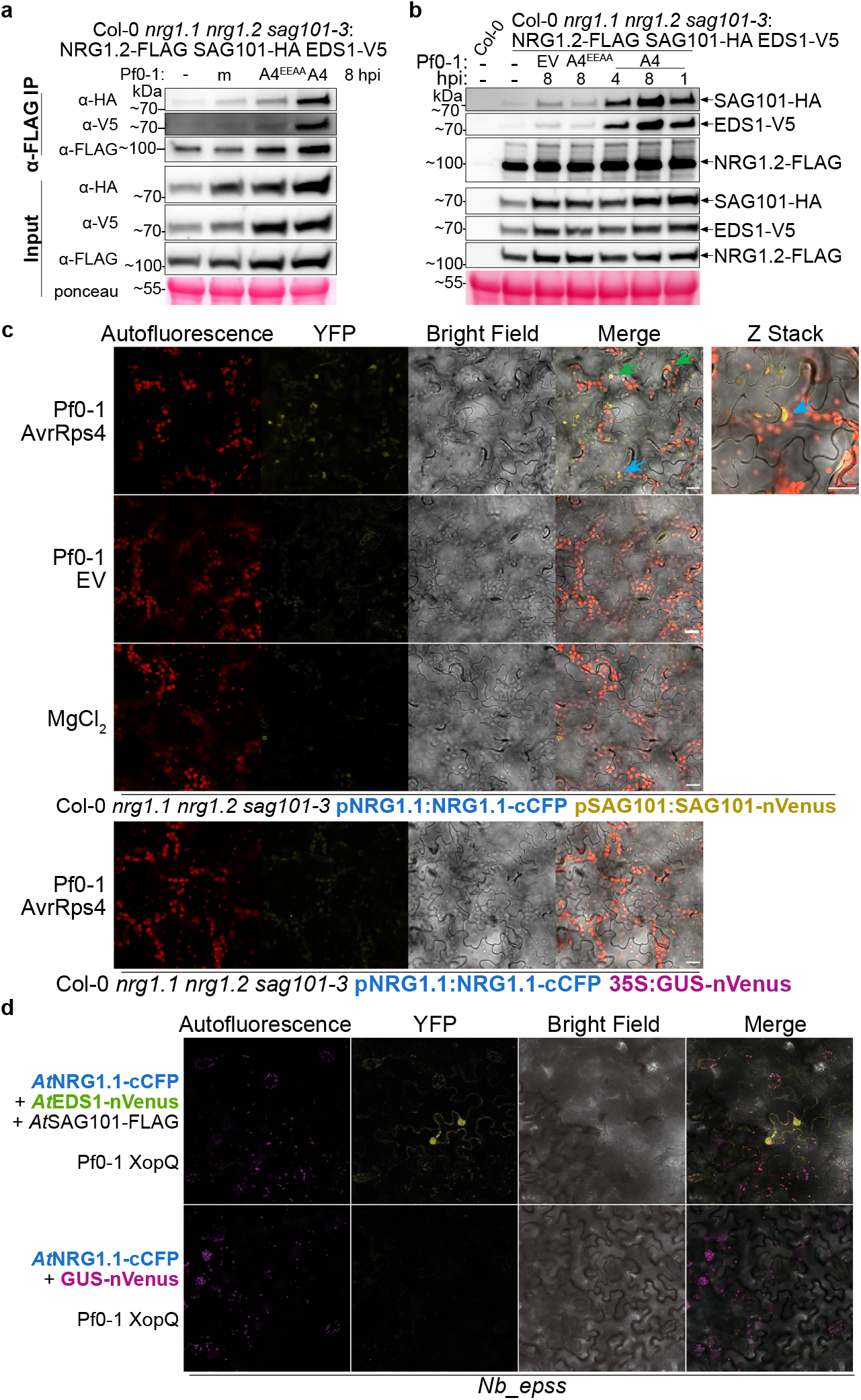
NRG1/EDS1 and NRG1/SAG101 associations at PM and in nucleus in Arabidopsis. **(a)** NRG1 weakly associates with EDS1 and SAG101 upon cell surface receptor activation in Arabidopsis. Experiments were performed on three biological replicates each for two independent lines, for a total of six replicates, with similar results. In (a) and (b) SDS-PAGE and Western blots of coIPs performed with native promoter-driven Arabidopsis stable line. “A4” indicates AvrRps4, “A4^EEAA^” indicates AvrRps4^EEAA^, “m” indicates MgCl_2_ mock, and “-” indicates un-infiltrated. **(b)** Effector delivery in Arabidopsis induces NRG1 interaction with EDS1 and SAG101. Experiments were performed on three biological replicates with similar results. **(c**) NRG1 associates with SAG101 in nuclei and at the PM post Pf0-1 AvrRps4 delivery in Arabidopsis. Live cell imaging of native promoter-driven Arabidopsis stable lines. Micrographs were taken 4-5 hpi of Pf0-1 and all images show single planes, excluding Z-stack image with Pf0-1 AvrRps4 treatment. Three biological replicates were performed with similar results, and representative micrographs are shown. Scale bar = 10 μM. Arrows indicate nuclei. Blue arrows indicate same nucleus between images. Autofluorescence (red) and YFP BiFC signal (yellow) are shown. (**d)** NRG1 associates with EDS1 in nuclei and at the PM post Pf0-1 XopQ delivery in *N. benthamiana. Agrobacteria* carrying 35S promoter-driven BiFC constructs were infiltrated into *Nb_epss* leaves. At 48 hpi, Pf0-1 XopQ was infiltrated. At 4-6 hpi, leaf disks were imaged. Three independent biological replicates, each with four technical replicates, were performed with similar results and representative images are shown. Autofluorescence (magenta) and YFP BiFC signal (yellow) are shown. Quantification of PM and nuclear signal in *SI Appendix*, Fig. 2b & 2c.

We investigated the subcellular locations of the induced EDS1 and SAG101 complexes with NRG1 upon effector recognition utilizing bimolecular fluorescence complementation (BiFC) assays. We complemented Col-0 *nrg1*.*1 nrg1*.*2 sag101-3* with pNRG1.1:NRG1.1-cCFP and pSAG101:SAG101-nVenus (*SI Appendix*, Fig. S1c) for NRG1-SAG101 BiFC assays. NRG1-EDS1 BiFC assays were carried out via Agro-mediated transient expression of 35S promoter-driven *At*NRG1.1-cCFP with *At*EDS1-nVenus and *At*SAG101-Flag in *N. benthamiana* (*SI Appendix*, Fig. S2a). Detection of YFP signal in BiFC assays indicates components are within sufficient proximity to reconstitute fluorophore excitation. An enhanced YFP signal was detected upon Pf0-1 AvrRps4 delivery in Arabidopsis (Fig. 1c) and also upon Pf0-1 XopQ delivery in *N. benthamiana* (Fig. 1d & *SI Appendix*, Fig. S2b & S2c). XopQ is recognized by the endogenous TNL Roq1 in *N. benthamiana*. In both assays, YFP signal was detected in the nucleus and at the PM. These data indicate that NRG1-SAG101 and NRG1-EDS1 associate in multiple locations within the cell.

Few data were reported that unambiguously support a nuclear-localized mechanism for NRG1 in defense responses (12, 16, 17). Therefore, we independently verified nuclear localization of NRG1 utilizing Pf0-1 delivery of AvrRps4 in Col-0 *nrg1*.*1 nrg1*.*2* carrying pNRG1.1:NRG1.1-mRuby (*SI Appendix*, Fig. S3a). Upon Pf0-1 delivery of AvrRps4, an increased mRuby signal is detected in nuclei (*SI Appendix*, Fig. S3c). These data indicate that NRG1 either re-localizes to nuclei upon effector recognition or is present in low abundance prior to immune activation. Importantly, they suggest a previously unreported defense-dependent NRG1 association with EDS1 and SAG101 in the nucleus.

### NRG1 oligomerizes upon effector-dependent defense activation in Arabidopsis

We investigated self-association of NRG1 in Arabidopsis before and after immune activation in super-transformed lines of Ws-2 *nrg1*.*1 nrg1*.*2* pNRG1.2:NRG1.2-FLAG (26) carrying an additional epitope-tagged NRG1, pNRG1.2:NRG1.2-V5. NRG1 self-association was evaluated by α-FLAG IP of NRG1.2-FLAG followed by α-V5 immunodetection of NRG1.2-V5. We tested this association after Pf0-1 delivery of AvrRps4 or AvrRps4^EEAA^, with un-infiltrated leaves as a pre-activation control. We found that NRG1.2-V5 associated with NRG1.2-FLAG only after AvrRps4 delivery (Fig. 2a). These data indicate that NRG1 does not interact with itself in the pre-activation state, or after surface receptor activation, but only upon TNL immune activation.

**Figure 2.**
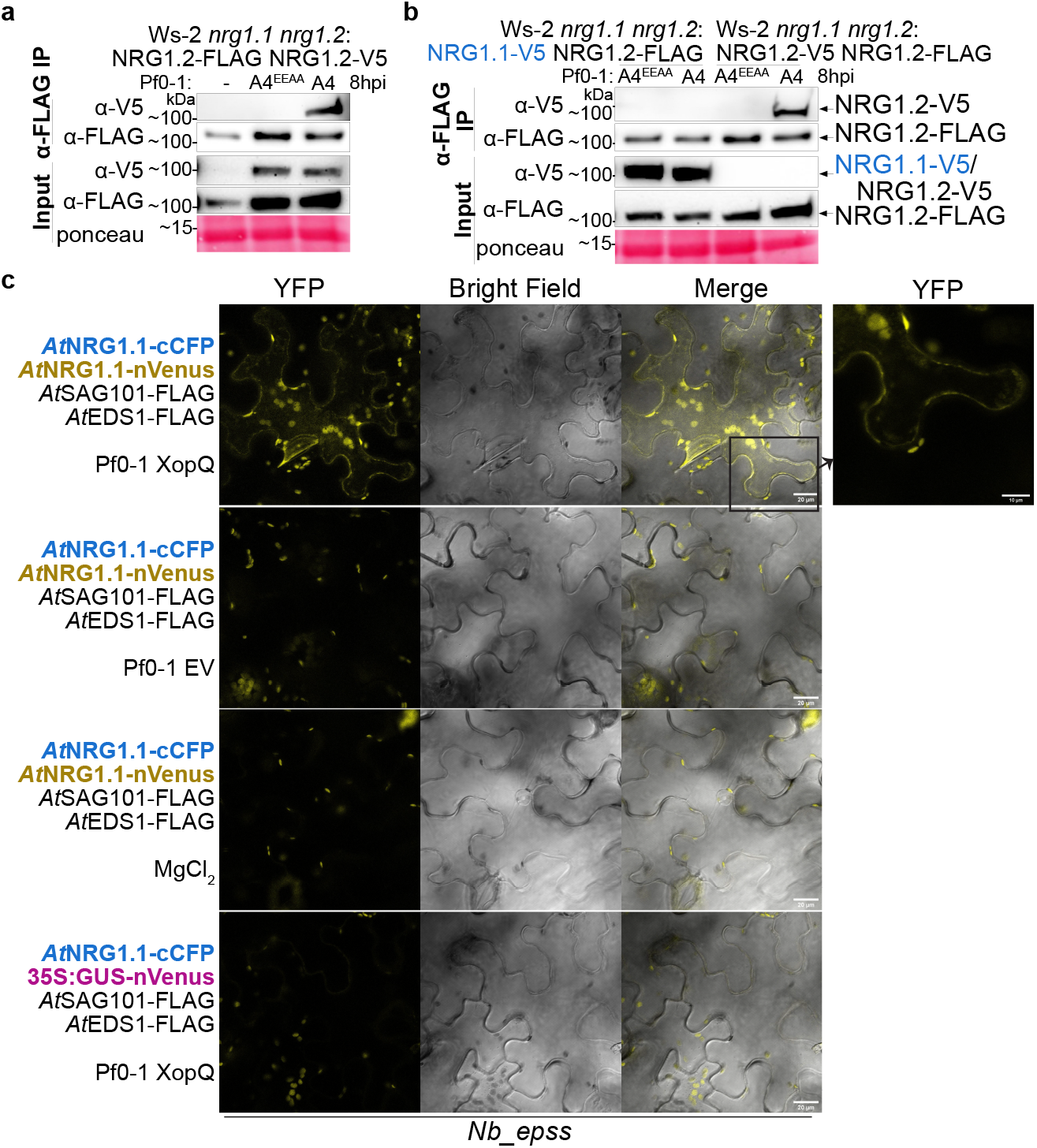
NRG1 self-associates upon AvrRps4 recognition in Arabidopsis. **(a)** NRG1.2-FLAG associates with NRG1.2-V5 upon AvrRps4 delivery in Arabidopsis. In (a) and (b), SDS-PAGE and Western blots of coIPs performed with native promoter-driven Arabidopsis stable lines. Experiments were performed on three biological replicates each for two independent lines, for a total of six replicates, with similar results. “A4” indicates AvrRps4, “A4^EEAA^” indicates AvrRps4^EEAA^, and “-” indicates un-infiltrated. **(b)** NRG1 self-association is paralog specific. **(c)** NRG1 self-association is localized to the PM. *Agrobacteria* carrying 35S promoter-driven BiFC constructs were infiltrated into *Nb_epss* leaves. At 48 hpi, Pf0-1 XopQ was infiltrated. At 8 hpi, leaf disks were imaged. Three independent biological replicates, each with four technical replicates, were performed with similar results and representative images are shown. Scale bar = 20 μM.

We also investigated whether NRG1 self-association is paralog specific. We super-transformed the previously generated Ws-2 *nrg1*.*1 nrg1*.*2* pNRG1.1:NRG1.1-V5 (26) to carry an additional variant of pNRG1.2:NRG1.2-FLAG. Unlike NRG1.2-V5, NRG1.1-V5 does not associate with NRG1.2-FLAG upon delivery of AvrRps4 (Fig. 2b). These data indicate that NRG1 self-association is paralog specific.

To determine the subcellular location of self-associating NRG1, we carried out *At*NRG1.1-cCFP and *At*NRG1.1-nVenus BiFC assays by Agro-mediated transient expression in *N. benthamiana* (*SI Appendix*, Fig. S2d). In three of five replicates, an enhanced YFP signal was observed only with Pf0-1 XopQ treatment (Fig. 2c). However, in two other replicates, a YFP signal was detected in control treatments, possibly due to overexpression or tag-specific auto-activity of NRG1 (11). We interpret these collective data as independent confirmation of the effector-dependent NRG1 self-association. Notably, in contrast to the NRG1-EDS1 and NRG1-SAG101 associations observed in nuclei and at the PM (Fig. 1c & 1d), NRG1-NRG1 YFP signal was only detected at the PM (Fig. 2c). These data indicate that while NRG1 forms associations with EDS1 and SAG101 in multiple compartments, NRG1 self-association is restricted to the PM.

Expression of auto-active *At*NRG1.1^D485V^ in *N. benthamiana* shows slower migrating forms in Blue Native PAGE assays (BN-PAGE) (Jacob et al., 2021) that may indicate NRG1 oligomers. To investigate NRG1 oligomer formation in Arabidopsis, we subjected lysates and elution products from Ws-2 *nrg1*.*1 nrg1*.*2* pNRG1.2:NRG1.2-FLAG pNRG1.2:NRG1.2-V5 to BN-PAGE assays. Species of NRG1.2-FLAG or NRG1.2-V5 were immunolabelled after α-FLAG IP of NRG1.2-FLAG. Slower migrating forms of NRG1.2-FLAG and NRG1.2-V5 were not observed in lysates after Pf0-1 AvrRps4 delivery in Arabidopsis. However, a slower migrating form of NRG1.2-V5 was observed in Pf0-1 AvrRps4-treated tissues after α-FLAG IP of NRG1.2-FLAG (Fig. 3a). Notably, no slow-migrating NRG1.2-FLAG was detected after α-FLAG IP of NRG1.2-FLAG (Fig. 3a) in Pf0-1 AvrRps4-treated tissue. These data indicate the formation of an effector-dependent NRG1 oligomer in Arabidopsis in a small proportion of total NRG1 protein after immune activation.

**Figure 3.**
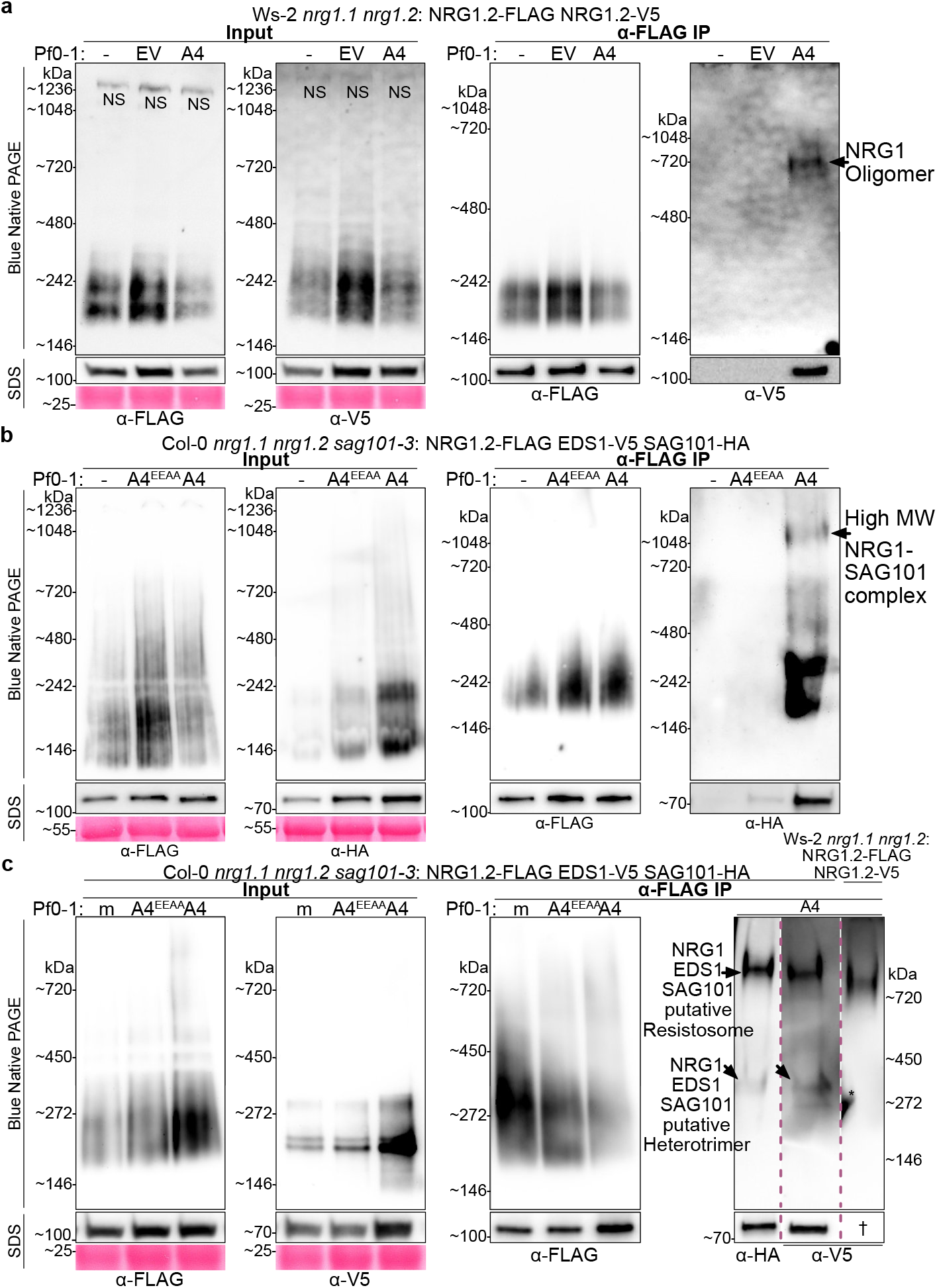
Formation of NRG1 oligomer in complex with EDS1 and SAG101 requires effector recognition in Arabidopsis. BN-PAGE and Western blot of lysates and coIP elution products performed with native promoter-driven Arabidopsis stable lines. “A4” indicates AvrRps4, “A4^EEAA^” indicates AvrRps4^EEAA^, “-” indicates un-infiltrated, and “m” is MgCl_2_ mock infiltration. Tissue was harvested 8 hpi. Experiments were performed on three biological replicates with similar results. “NS” indicates non-specific band. **(a)** Effector-dependent formation of an NRG1 oligomer in Arabidopsis. **(b)** Higher-order forms of NRG1-SAG101 complexes require effector delivery in Arabidopsis. **(c)** NRG1 oligomer co-migrates with the higher-order form of NRG1-associated EDS1-SAG101 in Arabidopsis after effector delivery. Red dash lines in BN-PAGE indicate lanes have been cropped from one membrane where samples were run in parallel. Red dash lines in SDS-PAGE indicate lanes have been cropped. Asterisk (**\***) indicates signal is bleed through from adjacent lane not shown. Dagger “†” indicates image already shown in separate figure.

### Effector-dependent association of EDS1 and SAG101 with an NRG1 oligomer in Arabidopsis

We investigated whether EDS1 and SAG101 interact with the NRG1 oligomer in an NRG1-EDS1-SAG101 Resistosome-like complex. We subjected pre- and post-activation lysates and elution samples from Col-0 *nrg1*.*1 nrg1*.*2 sag101-3* pNRG1.2:NRG1.2-FLAG pSAG101:SAG101-HA pEDS1:EDS1-V5 to BN-PAGE assays. As with NRG1.2 (Fig. 3a), slower migrating forms of SAG101-HA and EDS1-V5 were not observed in lysates after Pf0-1 AvrRps4 delivery in Arabidopsis (Fig. 3b, Fig. 3c). However, multiple forms of SAG101-HA were detected in BN-PAGE after α-FLAG IP of NRG1.2-FLAG in Pf0-1 AvrRps4 treated tissues, including a slow-migrating species above the 720 kDa marker (Fig. 3b). This contrasts with the presence of a single NRG1.2-V5 species after α-FLAG IP of NRG1.2-FLAG (Fig. 3a). Although SAG101-HA and EDS1-V5 weakly associated with NRG1.2-FLAG in pre-activation states (Fig. 1a & 1b), no SAG101-HA signal was detected after α-FLAG IP of NRG1.2-FLAG in un-infiltrated or Pf0-1 AvrRps4^EEAA^ infiltrated tissue (Fig. 3b), suggesting any pre-activation associations are of too low abundance to be detected. These data indicate formation of multiple NRG1-EDS1-SAG101 oligomeric complexes, in contrast to the single NRG1-NRG1 oligomer (Fig. 3a), upon immune activation in Arabidopsis.

Heterodimers of EDS1-SAG101 were previously found to elute near the 158 kDa marker in size exclusion chromatography (18) and a putative EDS1-SAG101 heterodimer migrates between 146 and 242 kDa markers in BN-PAGE assays (S*I Appendix*, Fig. S7). A species of SAG101-HA and EDS1-V5 is detected after α-FLAG IP of NRG1.2-FLAG in AvrRps4-treated tissues migrating just above the ∼272 kDa marker (Fig. 3b & 3c). It is possible these data represent degradation of NRG1-EDS1-SAG101 complexes into NRG1 monomers and EDS1-SAG101 heterodimers. However, an effector-dependent, low molecular weight species of SAG101-HA still migrates slower than the pre-activation species of SAG101-HA, EDS1-V5, or NRG1.2-FLAG (S*I Appendix*, Fig. S7). This activated complex contains NRG1.2-FLAG as it was isolated by α-FLAG IP. Yet, an NRG1.2-FLAG species is not detected near the ∼272 kDa marker and only the oligomeric species is detected above the ∼720 kDa marker (Fig. 3a & 3c). We interpret these collective data to reflect formation of a putative NRG1-EDS1-SAG101 heterotrimer (NRG1 monomer in complex with an EDS1-SAG101 heterodimer) upon effector recognition in Arabidopsis.

To investigate whether the slower migrating forms of NRG1, SAG101, and EDS1 correspond to the same molecular complex, elution products after α-FLAG IP of NRG1.2-FLAG from either Col-0 *nrg1*.*1 nrg1*.*2 sag101-3* pNRG1.2:NRG1.2-FLAG pSAG101:SAG101-HA pEDS1:EDS1-V5 or Ws-2 *nrg1*.*1 nrg1*.*2* pNRG1.2:NRG1.2-FLAG pNRG1.2:NRG1.2-V5 were resolved by BN-PAGE in parallel. Immunolabelling with α-HA for SAG101-HA and α-V5 for EDS1-V5 or NRG1.2-V5 revealed that AvrRps4-induced slow migrating forms of SAG101-HA, EDS1-V5, and NRG1.2-V5 after α-FLAG IP of NRG1.2-FLAG migrate at an indistinguishable size—consistent with an NRG1 oligomer in complex with EDS1 and SAG101 (Fig. 3c). These data indicate that effector recognition in Arabidopsis leads to accumulation of a putative NRG1-EDS1-SAG101 Resistosome.

### Effector recognition is sufficient to induce NRG1/EDS1 and NRG1/SAG101 association and requires TIR-domain enzyme activity

Plant TIR domains are holoenzymes (3–5, 27) which can produce a suite of cyclic and non-cyclic small molecules using NAD+ as an initial substrate. Products include nicotinamide and several ADPR-like molecules such as di-ADPR or ADPR-ATP that activate EDS1-dependent defense but which are insufficient for cell death (15, 27–30). We investigated whether TIR enzymatic activity is required for EDS1 and SAG101 association with NRG1 using Agro-infiltration in *N. benthamiana*. Expression of TIR-only Arabidopsis RBA1 is sufficient for ADR1-induced association with EDS1-family proteins in *N. benthamiana* (31). Utilizing a cell death inactive NRG1.2^EEAA^ mutant (S*I Appendix*, Fig. S4) which maintains TNL-induced association with EDS1 and SAG101 (12), we tested whether NRG1 could associate with EDS1 and SAG101 in the presence of RBA1 or NADase mutant RBA1^E86A^. We found that EDS1-V5 and SAG101-Myc associate with NRG1.2^EEAA^-FLAG when co-expressed with RBA1, but not with enzymatically inactive RBA1^E86A^ (S*I Appendix*, Fig. S5). These data indicate that TIR domain enzymatic activity is required for EDS1 and SAG101 association with NRG1 *in vivo*.

Macroscopic cell death in Arabidopsis, indicated by tissue collapse, requires both cell-surface receptor activation (PAMP-triggered immunity or “PTI”) and effector recognition by intracellular receptors (effector-triggered immunity or “ETI”) (20, 21). We investigated whether ETI in the absence of PTI is sufficient for NRG1 association with EDS1 and SAG101. We generated a Col-0 *nrg1*.*1 nrg1*.*2 sag101-3* line carrying pNRG1.2:NRG1.2-FLAG pSAG101:SAG101-HA pEDS1:EDS1-V5 with a β-estradiol-inducible AvrRps4-mCherry driven by the LexA promoter with the XVE system (32) (*SI Appendix*, Fig. S6a). Leaves were infiltrated with β-estradiol to induce AvrRps4-mCherry expression to activate ETI alone, or co-infiltrated with β-estradiol and Pf0-1 AvrRps4^EEAA^ for reconstitution of PTI+ETI co-activation. As previously, β-estradiol induction of AvrRps4 (ETI) is sufficient for defense gene activation (20) and induces weak microscopic cell death without tissue collapse, while β-estradiol induction of AvrRps4-mCherry in the presence of Pf0-1 AvrRps4^EEAA^ (PTI + ETI reconstitution) induces strong macroscopic cell death phenocopying Pf0-1 AvrRps4 delivery (PTI+ETI) (33) (S*I Appendix*, Fig. S6). We found that induction of AvrRps4-mCherry by β-estradiol infiltration is sufficient to induce EDS1-V5 and SAG101-HA association with NRG1.2-FLAG (Fig. 4a). Notably, the SAG101-HA and EDS1-V5 signal after α-FLAG IP of NRG1.2-FLAG is indistinguishable between β-estradiol (ETI), Pf0-1 AvrRps4 (PTI+ETI), and β-estradiol with Pf0-1 AvrRps4^EEAA^ (PTI+ETI reconstitution). These data indicate that ETI is sufficient to induce EDS1 and SAG101 interaction with NRG1 in Arabidopsis, and to a level comparable with PTI and ETI co-activation.

**Figure 4.**
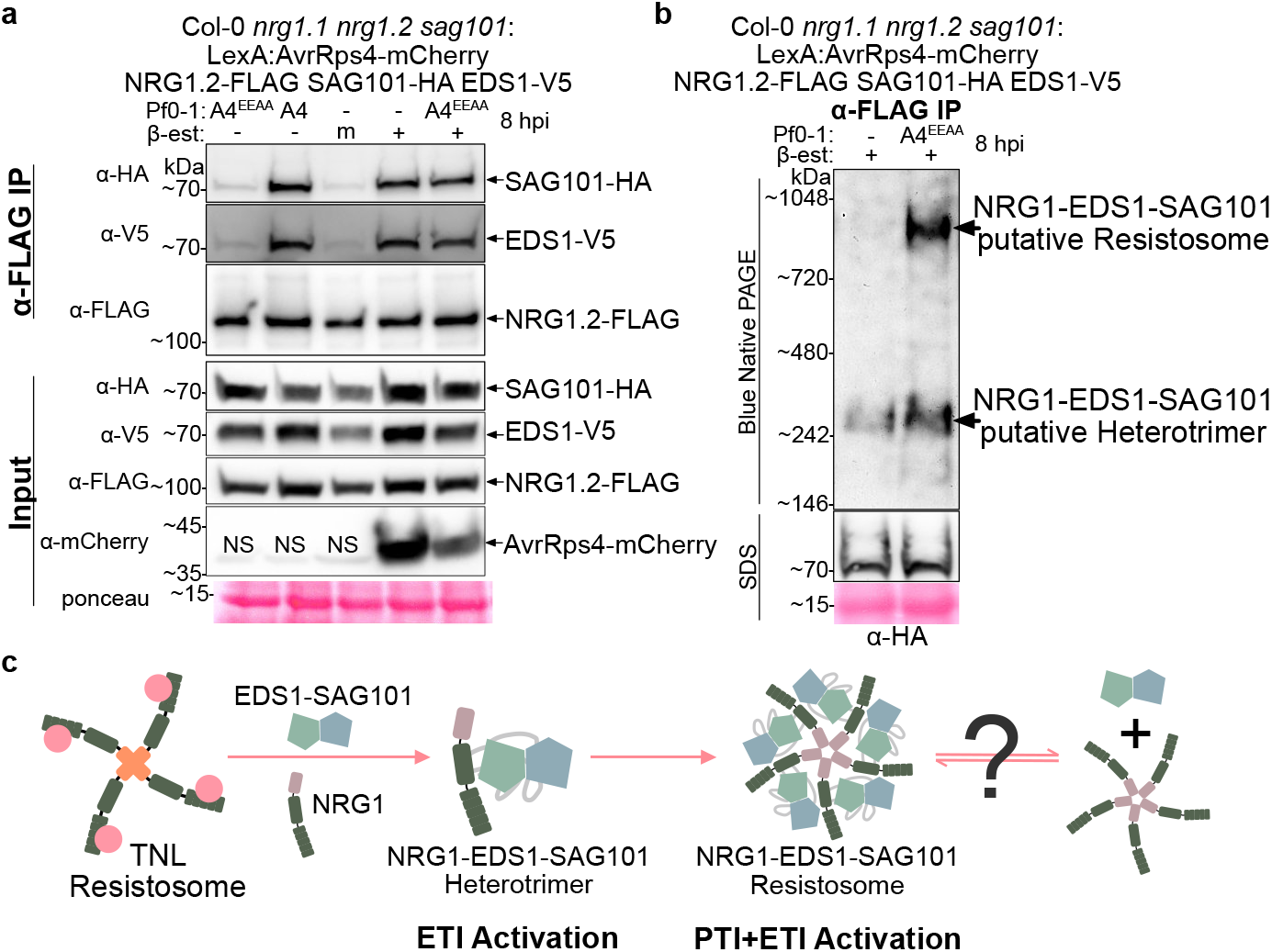
Both PTI and ETI activation are required for accumulation of NRG1-EDS1-SAG101 oligomeric “Resistosome” like complex. **(a)** ETI alone is sufficient to induce NRG1/EDS1 and NRG1/SAG101 associations in Arabidopsis. SDS-PAGE and Western blot of lysates and coIP elution products from native promoter-driven Arabidopsis stable lines. Experiments were performed on three biological replicates each for two independent lines, for a total of six replicates, with similar results. “m” indicates mock infiltration, “NS” indicates non-specific band, and “-” indicates no infiltration in (a) and (b). **(b)** Both ETI and PTI activation are required for accumulation of NRG1-EDS1-SAG101 complexes. BN-PAGE and Western blot of coIP elution products from native promoter-driven Arabidopsis stable lines. Experiments were performed on three biological replicates with similar results. **(c)** Schematic for proposed activation and oligomerization of NRG1 with EDS1 and SAG101. Upon TNL binding of effector ligands, ETI is activated and NRG1 is induced to associate with EDS1 and SAG101 to form an NRG1-EDS1-SAG101 heterotrimer. When PTI and ETI are co-activated, NRG1-EDS1-SAG101 Resistosomes are formed, potentially followed by dissociation of EDS1-SAG101 heterodimers to form a final NRG1 oligomer.

### PTI and ETI co-activation promote accumulation of the NRG1-EDS1-SAG101 oligomeric complex

To test if β-estradiol-induction of NRG1-EDS1-SAG101 complexes is sufficient for formation of NRG1-EDS1-SAG101 Resistosomes, we performed BN-PAGE with elution products from Col-0 *nrg1*.*1 nrg1*.*2 sag101-3* carrying pNRG1.2:NRG1.2-FLAG pSAG101:SAG101-HA pEDS1:EDS1-V5 pLexA:AvrRps4-mCherry XVE after β-estradiol induction of AvrRps4-mCherry (ETI) or β-estradiol co-infiltration with Pf0-1 AvrRps4^EEAA^ (PTI+ETI reconstitution). As expected, β-estradiol co-infiltration with Pf0-1 AvrRps4^EEAA^ induces the accumulation of a slow-migrating SAG101-HA species, above the 720 kDa marker, after α-FLAG IP of NRG1.2-FLAG (Fig. 4b). However, this high molecular weight species is not detected after β-estradiol induction of AvrRps4-mCherry, and only the low molecular weight SAG101-HA species above the 242 kDa marker is present (Fig. 4b). Although ETI is sufficient for NRG1 interaction with EDS1 and SAG101, these data indicate ETI is not sufficient for accumulation of high molecular weight NRG1-EDS1-SAG101 complexes. This finding highlights a requirement for an unknown PTI activation-dependent mechanism facilitating *in vivo* assembly of a putative NRG1-EDS1-SAG101 Resistosome.

## Discussion

Co-activation and mutual potentiation of cell surface-receptor (PTI) and intracellular receptor (ETI) immune systems are required to mount a pathogen-restricting defense and host cell death response in plants (20, 21) but the mechanism by which PTI enhances cell death upon ETI is unknown. We show here that while ETI alone is sufficient for NRG1 association with EDS1 and SAG101 (Fig. 4a), ETI is not sufficient for formation of the NRG1-EDS1-SAG101 putative Resistosome (Fig. 4b). Formation of higher-order NRG1 complexes was reported to correlate with activation of cell death (16). Our data suggest that PTI is required for TNL-dependent cell death because it potentiates formation of a high molecular weight NRG1-EDS1-SAG101 Resistosome-like complex.

The helper NLR NRG1 is essential for TNL-mediated cell death via a mechanism that involves EDS1 and SAG101 (10–13, 17, 26). A related helper NLR, ADR1, functions together with EDS1 and PAD4 for transcriptional amplification of defense genes (12, 13, 19, 22, 31). In Arabidopsis, these modules appear to partially compensate for the other’s absence (12, 13). This indicates these complexes contribute to Arabidopsis TNL-triggered defense responses to different extents. We speculate that TNL-triggered cell death and transcription mediated by NRG1-EDS1-SAG101 are carried out by distinct complexes that are spatially separated within the cell: Resistosomes at the PM and heterotrimers in the nucleus. We confirmed the requirement for PTI and ETI co-activation in macroscopic cell death (33) (S*I Appendix*, Fig. S6). We find that NRG1 self-associates primarily or exclusively at the PM (Fig. 2c) but interacts with EDS1 and SAG101 at both the PM and in nuclei (Fig. 1c & 1d). As the putative NRG1-EDS1-SAG101 Resistosome is formed under PTI and ETI co-activation (Fig. 4b), it is likely that this form contributes to cell death at the PM. Indeed, previous reports identify a cell death mechanism for an oligomerized, constitutively active NRG1 at the PM (16). We also confirmed the sufficiency of ETI activation for transcription elevation of defense genes (20) (S*I Appendix*, Fig. S6). The lack of an oligomerized NRG1 in the nucleus or formation of a putative NRG1-EDS1-SAG101 Resistosome during ETI alone (Fig. 4b), points to the presence of NRG1-EDS1-SAG101 heterotrimers mediating transcriptional defense responses in the nucleus. Our inference that NRG1 has a nuclear activity is consistent with earlier studies showing that nuclear-enriched EDS1 restricted pathogen growth in Arabidopsis (34, 35). How heterotrimer immune complexes in nuclei, containing non-oligomerized helper NLRs, might function in transcriptional defense remains unclear.

We show that TIR domain NADase activity is required for NRG1 association with EDS1 and SAG101 (S*I Appendix*, Fig. S5). Indeed, TNL-mediated ETI activation leads to interaction of NRG1-EDS1-SAG101 (15) and ADR1-EDS1-PAD4 (14) complexes via TIR domain catalysis of distinct small molecules. Likely, the production and availability of TIR small molecules dictate activation of NRG1-EDS1-SAG101 versus ADR1-EDS1-PAD4 signalling pathways. Small-molecule-induced NRG1-EDS1-SAG101 heterotrimer formation is analogous to formation of ZAR1-RKS1 intermediate species upon interaction with uridylylated PBL2 (6). Conceivably, steric changes induced by NRG1-EDS1-SAG101 heterotrimer formation enable Resistosome assembly in a similar manner to ZAR-RKS1-PBL2^UMP^ Resistosomes (6, 7). However, in contrast to what is known for ZAR1, a PTI mechanism is also required for formation of NRG1-EDS1-SAG101 Resistosomes *in vivo*, as ETI alone is insufficient (Fig. 4b). Furthermore, a function for heterotrimers in nuclear signalling is not hypothesized for ZAR1, as studies only found high molecular weight ZAR1 oligomers which form plasma membrane pores (9). Our study highlights the novel possibility that oligomeric and heterotrimeric helper NLR forms may have distinct functions in mediating defense outputs in plants.

In contrast to previous studies which showed 100% conversion of NRG1.1 to slower-migrating species of auto-active NRG1.1^D485V^ in BN-PAGE (16), only a small proportion of NRG1.2, EDS1, and SAG101 protein was converted to putative Resistosomes in Arabidopsis (Fig. 3). This also contrasts with the quantitative conversion of inactive to oligomerized helper NLR NRC in Solanaceae (36, 37). It may be that very few activated forms of NRG1-EDS1-SAG101 Resistosomes are sufficient for defense, which potentially increase Ca^2+^ influx to the cytoplasm (15). However, it is plausible that low accumulation of high molecular weight complexes is an artifact of enrichment bias for pre-activated species during IP. Conceivably, either activated oligomeric species are outnumbered by pre-activation species or nuclear forms of NRG1, or previously exposed epitopes might become buried, or changes in localization inhibit extraction. Low detection of Resistosome accumulation could be further influenced by NRG1-EDS1-SAG101 oligomeric complex instability. Perhaps, EDS1-SAG101 promotes NRG1 oligomerization to then exit the complex, leaving a putative NRG1 pentamer (Fig. 4c). Indeed, faster migrating species of EDS1-SAG101 were detected after IP of NRG1 in activated tissues, including a species < 242 kDa and lower molecular weight than the putative heterotrimer (Fig. 3b, 3c, & 4b), which could be interpreted as dissociation from NRG1 oligomers. However, only one stable form of oligomeric NRG1.2 was observed *in vivo* (Fig. 3a & 3c) and migrated indistinguishably from the high molecular weight species of SAG101 and EDS1 after IP of NRG1.2 in activated tissues (Fig. 3c). This indicates that once putative NRG1-EDS1-SAG101 Resistosomes are formed, they are stable. This shares similarity with the ZAR1 pentamer which remains stably associated with RKS1/PBL2^ump^ in Resistosomes once formed (6, 7). Our data indicate the stable formation of an immune activation-dependent putative NRG1-EDS1-SAG101 Resistosome in Arabidopsis and highlight the importance of investigating NRG1 molecular assemblies and sub-cellular sites of action in the presence of EDS1 and SAG101.

The requirement for PTI and ETI co-activation in TNL-mediated cell death (33) is intriguing in light of our observation that PTI and ETI co-activation is required for detection of putative NRG1-EDS1-SAG101 Resistosomes (Fig. 4b). It should not be excluded that assembly of NRG1-EDS1-SAG101 Resistosomes could occur, below detection thresholds, under ETI activation alone. Conceivably, PTI-mediated modification(s) of NRG1, EDS1, and/or SAG101 stabilize Resistosomes, allowing accumulation at the PM to function in increasing cytoplasmic [Ca^2+^] and promotion of cell death. Alternatively, PTI could more indirectly facilitate accumulation of NRG1-EDS1-SAG101 Resistosomes *in vivo*. PTI activation leads to transcriptional accumulation of TIR-domain proteins (22), which may enhance EDS1-SAG101 assisted assembly and oligomerization of NRG1 Resistosomes through increased production of nucleotide-based small molecules. Our study highlights the importance of future investigations into the possible role of NRG1-EDS1-SAG101 complexes as a mechanistic link for PTI and ETI co-activation of plant defense responses.

## Materials and Methods

### NRG1.1 and NRG1.2

Arabidopsis carries functionally redundant paralogs NRG1.1 and NRG1.2 (17, 26), as well as the truncated NRG1.3 which is reported to antagonize NRG1-mediated immunity (38). This study employed either NRG1.1 or NRG1.2 to facilitate both biochemistry and cell biology investigations. When transiently expressed in *N. benthamiana, At*NRG1.1 shows an additional band in western blot assays (12) that may interfere with interpretations, while *At*NRG1.2 does not (*SI Appendix*, Fig S5). Furthermore, NRG1.2-FLAG was previously shown to be functional (26) and assays were optimized for α-FLAG IP. Although only one band is detected by western blot when NRG1.1 is expressed in Arabidopsis (Fig. 2b, *SI Appendix*, Fig S3), for simplicity between systems, NRG1.2 was preferred for biochemistry assays. Although both NRG1.1 and NRG1.2 are sensitive to fluorophore-tag specific defects (11), NRG1.1 was preferred in cell biology assays as loss of function or auto-active phenotypes were less likely.

### Growth of Arabidopsis & *N. benthamiana*

Arabidopsis plants for pathogen assays were grown with 8 h photoperiod in controlled environment rooms (CERs) at 20-22 °C with 70% relative humidity. Arabidopsis plants for seed collection were grown at similar CER conditions and a 16 h photoperiod. *N. benthamiana* plants for transient infiltration and cell death assays were grown with a 16 h photoperiod in a CER at a 20-22 °C with 70% relative humidity.

### Preparation of Pf0-1 for infiltration

The Effector-to-Host Analyzer (EtHAn) system (23) is a non-pathogenic strain of *Pseudomonas fluorescens* (Pf0-1) engineered with type III secretion system to deliver encoded effectors into plant cells. Glycerol stocks of Pf0-1 strains carrying effector expression-vectors were incubated 24-48 h at 28 °C on King’s B medium (KB) with antibiotics as previously described (23). Fresh colonies were cultured and incubated 24 h at 28 °C before suspension in MgCl_2_-MES pH 5.6.

### Pf0-1 infiltration of Arabidopsis leaves

Rosette leaves of Arabidopsis were infiltrated with Pf0-1 strains resuspended at OD_600_ = 0.2 in MgCl_2_-MES pH 5.6. Mock (“MgCl_2_”) infiltrations were MgCl_2_-MES pH 5.6 only. Leaves for protein assays were harvested, flash-frozen in liquid nitrogen, and stored at – 80 °C. Plants were ∼ 4-week-old in cell death complementation assays. Leaves were visualized for macroscopic tissue-collapse 24 h post effector-delivery by Pf0-1 infiltration. Plants were ∼ 4-5-week-old in coIP assays and siblings were visualized for HR 24 h post effector-delivery by Pf0-1 infiltration.

### β-estradiol infiltration of Arabidopsis leaves

Arabidopsis stable lines carrying the XVE cassette (32) were generated for β-estradiol induction of LexA promoter-driven AvrRps4-mCherry. Leaves of 4-5-week-old Arabidopsis were infiltrated with 50 μM β-estradiol diluted in MgCl_2_-MES pH 5.6 with 0.01% Silwet® L-77. Mock was prepared with dimethyl sulfoxide (DMSO) in place of β-estradiol.

### Agro-infiltration for cell death and co-immunoprecipitation assays

Glycerol stocks of *A. tumefaciens* strains were struck on LB agar plates containing antibiotic selections and incubated 48-72 h at 28 °C. Cells were collected from plates and resuspended in MgCl_2_-MES pH 5.6 at OD_600_ = 0.1-1. Silencing suppressor p19 (39) (OD_600_ = 0.5) and 200 μM acetosyringone were co-infiltrated into leaves of 4-6-week-old *N. benthamiana* leaves. In cell death assays, leaves were spot-infiltrated and evaluated for macroscopic tissue collapse 3-7 days post infiltration (dpi). In coIP assays, entire leaf surface was infiltrated and leaves were harvested 48-72 hpi, flash-frozen, and stored at – 80 °C.

### Localization and BiFC analysis with transgenic Arabidopsis

Leaves of 4-week-old transgenic Arabidopsis were infiltrated with (OD_600_ = 0.2) Pf0-1 AvrRps4, Pf0-1 EV, or 10 mM MgCl_2_ (mock) for live cell imaging. Leaf discs were harvested 4-6 h post Pf0-1 infiltration and imaged with a Zeiss LSM780 confocal laser scanning microscope for NRG1.1-cCFP and SAG101-nVenus BiFC, and with a Zeiss LSM700 confocal laser scanning microscope for NRG1.1-mRuby localization. The following excitation and detection windows were used: YFP 514 nm, 530-550 nm, mRuby 560 nm, 607-683 nm, autofluorescence of chlorophyll A 550 nm, 695-737 nm. Objectives used were: 20 x (0.8 NA, water), 40x (1.3 NA oil). DAPI staining was used to mark nuclei. Confocal images were compiled using ImageJ and Z-stacks were projected with Fiji using standard deviation methods.

### BiFC analysis in *N. benthamiana epss* reconstitution assays

*Agrobacterium* carrying BiFC constructs were syringe-infiltrated into leaves of *N. benthamiana* lacking all EDS1-family proteins (*Nb*_*epss*) (11) for live cell imaging. At 48 hpi of *Agrobacterium*, the leaf zone was subsequently infiltrated with (OD_600_ = 0.3) Pf0-1 XopQ, Pf0-1 EV, or 10 mM MgCl_2_ (mock). At 8 hpi of Pf0-1, leaf discs were harvested for confocal microscopy imaging. A Leica TCS SP8 confocal laser scanning microscope was used for NRG1.1-cCFP and EDS1-nVenus BiFC. A Zeiss LSM700 confocal laser scanning microscope was used for NRG1.1-cCFP and NRG1.1-nVenus BiFC. Excitation and detection windows used: YFP 514 nm, 530-550 nm, autofluorescence of chlorophyll A 550 nm, 695-737 nm. Objectives used: 20 x (0.8 NA, water), 40x (1.3 NA oil or 1.2 water). Confocal images were compiled using ImageJ and Z-stacks were projected with Fiji using standard deviation methods.

### Leaf protein accumulation assays

Leaf protein accumulation assays in *SI Appendix* Fig S3 were performed as described in (27).

### Arabidopsis lysate preparations

Protein was purified from ∼ 2.5-3.5 g dry weight Arabidopsis leaves flash-frozen in liquid nitrogen. Tissue was ground by mortar and pestle in liquid nitrogen and membranes were solubilized in 100 mM HEPES (pH 7.5), 300 mM NaCl, 5 mM MgCl_2_, 0.5% Nonidet^™^ P-40 Substitute (11754599001), 10% glycerol, 2% polyvinylpolypyrrolidone (PVPP), 1 tablet cOmplete^™^, EDTA-free protease inhibitor cocktail tablet, 10 mM DTT. Lysates were incubated inverting 10 min at 4 °C before centrifugation at 4 °C 35 min at 4000 x *g*. Lysates were filtered through Miracloth (475855). Input samples were combined with SDS sample buffer and 10 mM DTT and heated at 65 °C for 5 min before storage at – 20 °C.

### Co-immunoprecipitation with Arabidopsis lysates

Approx. 3-4 mL soluble lysate was combined with 50 μL Anti-FLAG^®^ M2 Affinity Gel slurry and inverted ∼ 2 h at 4 °C. Samples were centrifuged at 800 x *g* for 5 min at 4 °C, supernatant was removed, and beads were washed [100 mM HEPES (pH 7.5), 300 mM NaCl, 5 mM MgCl_2_, 0.5% Nonidet^™^ P-40 Substitute, 10% glycerol] inverting at 4 °C for 5 min for a total of 3 washes. Elution was performed in 100 μL wash buffer with 0.5 mg/mL 3×FLAG^®^ peptide. Sample was incubated ∼ 2.5 h at 4 °C shaking at 750 RPM for 5 min every 25 min. Elution product was transferred to fresh 1.5 mL Eppendorf. Beads were heated at 65 °C for 5 min in SDS sample buffer and 100 mM DTT. Elution product was combined with SDS sample buffer and 10 mM DTT, heated 65 °C for 5 min, and stored at – 20 °C.

### *N. benthamiana* lysate preparations

As described for Arabidopsis except that ∼ 0.5-1 g dry weight leaf tissue was used.

### Co-immunoprecipitation with *N. benthamiana* lysates

As described for Arabidopsis except that 1.5 mL lysate was combined with 20 μL Anti-FLAG^®^ M2 Affinity Gel slurry.

### Golden Gate Assembly of DNA constructs

Golden Gate cloning (40) was used for generation of stable Arabidopsis lines and transient expression in *N. benthamiana*. Arabidopsis lines for coIP were generated with native promoters and terminators with the exception of pSAG101:SAG101-HA:HSP18t. Arabidopsis lines for BiFC were generated with native promoters and terminators with the exception of pSAG101:SAG101-nVenus:35St. Transient expression constructs for coIP were generated with 35S promoters and Ocs terminators. Transient expression constructs for BiFC were generated with 35S promoters and 35S terminators and NRG1.1, EDS1, and SAG101 genomic sequences as described previously (12). C-terminal cCFP and nVenus tags were described previously (41). “FLAG” tag refers to 6×HIS-3×FLAG^®^ and “HA” refers to 6×HA. Golden Gate compatible epitope tag mRuby was synthesized by Thermo Fisher Scientific.

### SDS-PAGE

SDS sample buffer was prepared to a 4× concentration at 250 mM Tris-HCl (pH 6.8), 8% (w/v) sodium dodecyl sulphate (SDS), 40% glycerol (v/v), 50 mM EDTA, and bromophenol blue for visualization. Samples were prepared with SDS sample buffer to a final concentration of 1×, incubated 5 min at 65 °C, and stored at – 20 °C. Frozen samples were warmed to 37 °C for 5 min, briefly vortexed, and centrifuged at 15,000 x g for 1 min. Samples were resolved by SDS-PADE (precast 4-20% Mini-PROTEAN^®^ TGX^™^: 4561095) assembled in gel tank apparatus (Mini-PROTEAN^®^ system) with 1× SDS buffer [25 mM Tris, 200 mM Glycine, 2% SDS (w/v)]. Samples were loaded alongside 5 μL PageRuler^™^ Prestained Protein Ladder 10-180 kDa (26617). Electrophoresis was run at 90 V for ∼ 15 min then 135 V for ∼ 45 min or until dye-edge migrated to gel base. Input membranes were ponceau-stained as loading control.

### Blue Native PAGE

BN-PAGE was performed according to the NativePAGE^™^ Novex^®^ Bis-Tris Gel System with precast NativePAGE^™^ 3-12% Bis-Tris Mini Protein Gels (10-well BN1001BOX) and 5 μL NativeMark^™^ Unstained Protein Standard (LC0725), or SERVA (39219.01) marker in Fig. 3c only. Electrophoresis was run at 150 V for ∼ 60 min then at 250 V for ∼ 45 min until dye-edge migrated to gel base. Electrophoresis was run with dark cathode buffer for ∼25 min and then with light cathode buffer the remainder of the run. The gel was subjected to semi-dry transfer (described below). Membrane was incubated for 15 min in 8% acetic acid, rinsed, air-dried at RT > 1 h. Markers were visualized when membrane was pre-activated with ethanol. Figures were generated with SDS-PAGE and Western blot analysis of identical sample.

### Semi-dry protein transfer & Western blotting

Semi-dry transfer was performed using Bio-Rad Trans-Blot® Turbo^™^ Transfer System (1620177; Midi 0.2 μM PVDF Transfer Kit: 1704273) in standard settings. PVDF membranes were blocked 30-60 min, rotating 60 RPM at RT, with 5% milk TBS-T (50 mM Tris-HCl pH 8.0, 150 mM NaCl, 0.1% Tween^®^-20) before immunolabelling with α-V5-horseradish peroxidase (HRP), α-HA-HRP, α-FLAG^®^-HRP, or α-Myc-HRP in 5% milk TBS-T overnight at 4 °C or > 1 h at RT rotating 60 RPM. AvrRps4-mCherry and NRG1.1-mRuby were immunolabelled with α-mCherry, washed and re-probed with α-rabbit (Rb)-HRP. Membranes were washed three times for 5-10 min with TBS-T then three times 5-10 min with TBS rotating RT at 60 RPM. Membranes were incubated with SuperSignal^™^ West Pico PLUS (34580) or West Femto (34095) Chemiluminescent Substrate and imaged with GE Healthcare ImageQuant^™^ LAS 4000 or Amersham ImageQuant^™^ 800 enhanced chemiluminescence systems.

## Acknowledgments

We thank Tim Wells with JIC horticulture services for care of Arabidopsis and *N. benthamiana* plants, Mark Youles and Liam Egan with TSL Synbio, Ola Wawryk, Jodie Taylor, Jaqueline Bautor and Matthew Smoker for transformation of Arabidopsis, Johannes Stuttmann and Ulla Bonas for Pf0-1 XopQ, Marc Nishimura and Adam Bayless for RBA1 constructs, Brian Staskawicz for *Agrobacterium* XopQ, Dan Mclean for guidance generating Derevnina plots, Ruby O’Grady for illustrations, and Ton Timmers at MPI-Cologne for confocal microscopy advice. Work in the Parker lab was supported by the Max-Planck Society, Deutsche Forschungsgemeinschaft (DFG) grant SFB-1403–414786233 (J.E.P., X.S.) and an Alexander von Humboldt Foundation postdoctoral fellowship (J.W.). Work in the Jones lab was supported by ERC Advanced Investigator grant “ImmunityByPairDesign” (H-K.A.), BBSRC DTP program (B.P.M.N.) and by Gatsby Foundation support to TSL (J.M.F. and J.D.G.J.).

## Supplementary Figures

**Fig. S1.**
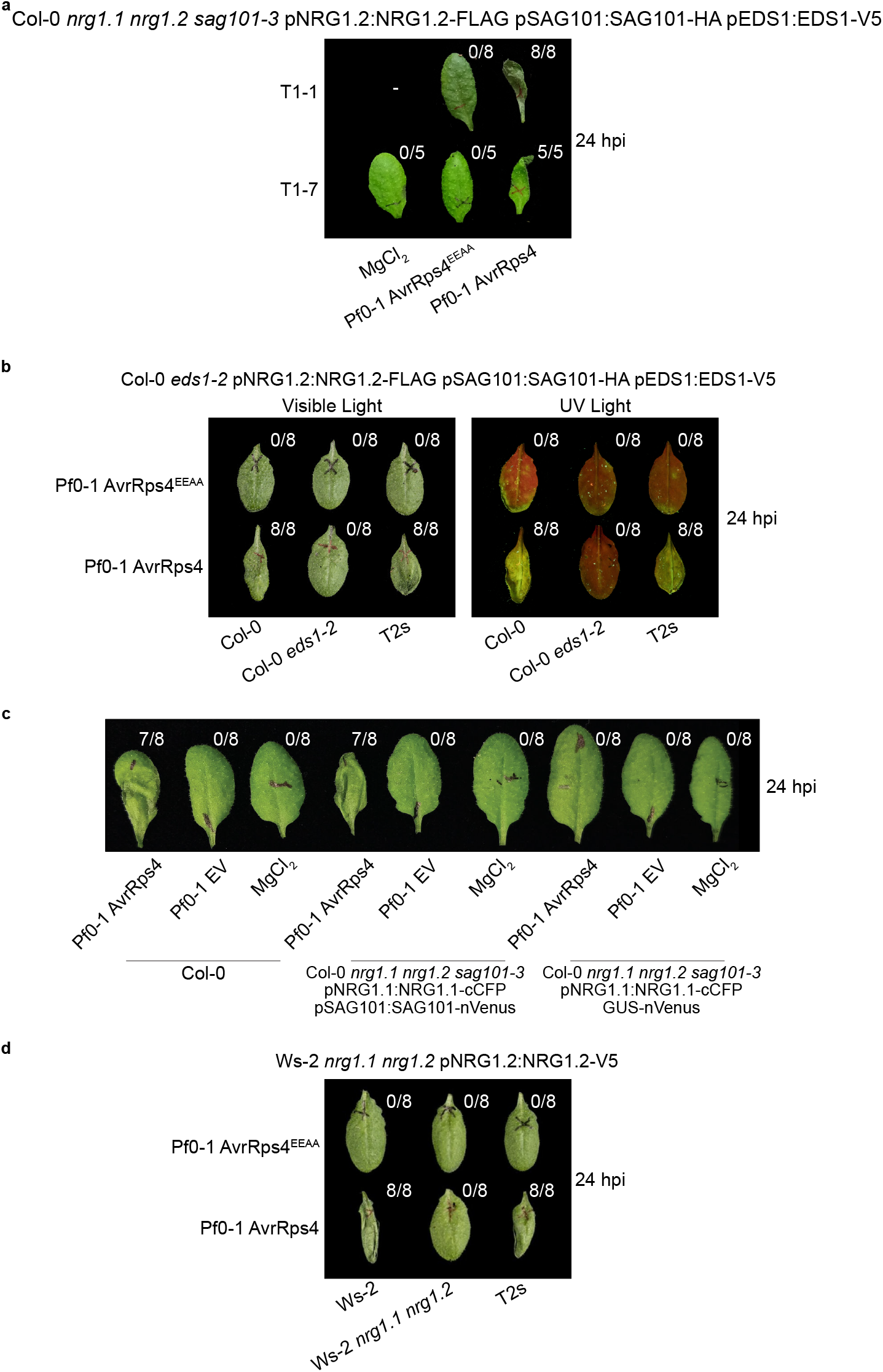
Cell death complementation in Arabidopsis stable transgenic lines. (a-d) Numbers in parentheses indicate number of leaves showing visual tissue collapse/total number of infiltrated leaves per genotype. Images were collected 24 hpi and a representative biological replicate is shown. **(a)** Stable expression of pNRG1.2:NRG1.2-FLAG and pSAG101:SAG101-HA complement Col-0 *nrg1*.*1 nrg1*.*2 sag101-3*. Pf0-1 AvrRps4 induces macroscopic cell death in leaves of two independent Arabidopsis lines. Pf0-1 AvrRps4^EEAA^ and MgCl_2_ are used as negative controls. Similar results were observed in more than three independent biological replicates. **(b)** Stable expression of pEDS1:EDS1-V5 complements Col-0 *eds1-2*. This was performed to confirm functionality of pEDS1:EDS1-V5 in (a) lines. Pf0-1 AvrRps4 induces macroscopic cell death in leaves of eight independent T2 generation individuals. Col-0 is used as a positive control, and Col-0 *eds1-2* is used as a negative control, for AvrRps4 recognition. Pf0-1 AvrRps4^EEAA^ is used as a negative control for cell death. White light images were collected to demonstrate tissue collapse. Ultraviolet light images were collected to demonstrate cell leakage. **(c)** Stable expression of pNRG1.1:NRG1.1-cCFP and pSAG101:SAG101-nVenus complement Col-0 *nrg1*.*1 nrg1*.*2 sag101-3* while 35S:GUS-nVenus does not. Col-0 was used as a positive control for AvrRps4 recognition. Pf0-1 EV and MgCl_2_ were used as negative controls for cell death. Three biological replicates were performed in one stable line with similar results. **(d)** Stable expression of pNRG1.2:NRG1.2-V5 complements Ws-2 *nrg1*.*1 nrg1*.*2*. This was performed to confirm functionality pNRG1.2:NRG1.2-V5 in super-transformed lines (Fig. 2). Pf0-1 AvrRps4 induces macroscopic cell death in leaves of eight independent T2 generation individuals. Ws-2 was used as a positive control and Ws-2 *nrg1*.*1 nrg1*.*2* was used as a negative control for AvrRps4 recognition. Pf0-1 AvrRps4^EEAA^ is used as a negative control for cell death.

**Fig. S2.**
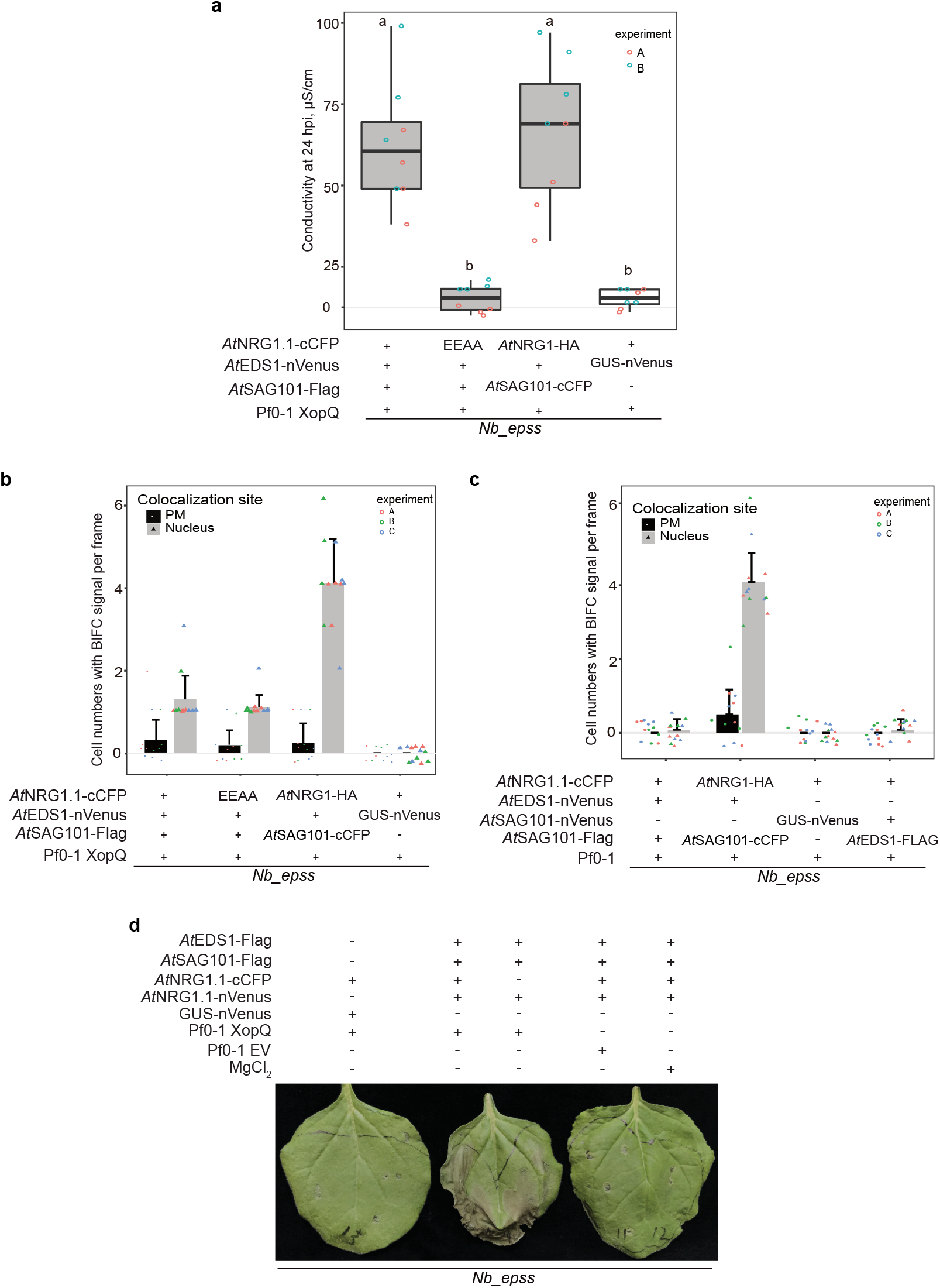
Quantification of YFP signal in *N. benthamiana* transient BiFC assays. **(a)** Constructs used in Fig. 1d and *SI Appendix*, Fig. S2c restore XopQ-triggered cell death in *Nb*_*epss* mutant background. Cell death was quantified in electrolyte leakage assays 6 h after harvesting leaf discs (24 hpi of Pf0-1 XopQ). Leaves were Agro-infiltrated 48 h prior to Pf0-1 infiltration with expression vectors carrying 35S promoter-driven *At*NRG1.1-cCFP or *At*NRG1.1^EEAA^-cCFP, *At*EDS1-nVenus or GUS-nVenus, and *At*SAG101-FLAG. Cell death was not observed with mutant *At*NRG1.1^EEAA^-cCFP or negative control GUS-nVenus. **(b)** Barplot quantification of YFP signal in Fig. 1c BiFC assays shows NRG1 co-localizes with EDS1 to PM and nuclei upon effector delivery. Co-localizations are not abolished with NRG1.1^EEAA^ mutant, which maintains association with EDS1 and SAG101 (12). NRG1^EEAA^ mutant is predicted to contact PM yet not form functional pore/channel (7, 12). EDS1 and SAG101 also co-localize to PM and nuclei, with a larger pool in nuclei. The number of cells with nuclear or PM YFP signal per frame were counted. **(c)** Barplot quantification of YFP signal in BiFC assays shows NRG1 does not co-localize with EDS1 upon cell-surface activation in the absence of effector delivery. EDS1 and SAG101 maintain their co-localization to nuclei and PM, with a larger pool in nuclei. The number of cells with nuclear or PM YFP signal per frame were counted. Experiment in (a) was performed two times independently, experiments in (b) and (c) were performed three times independently, each with four leaf-disc replicates. (Tukey’sHSD, α=0.001, n=8). Error bars represent standard error of mean. Datapoints with same colour come from one independent experiment. Boxplots and barplots were generated with ggplot2 (3.3.2) package in R. “EEAA” refers to *At*NRG1.1^EEAA^-cCFP. **(d)** Constructs used in NRG1.1 self-association assay in Fig. 2c rescue XopQ-triggered cell death when transiently expressed in *Nb*_*epss*. Leaves were Agro-infiltrated 48 h prior to Pf0-1 XopQ infiltration with 35S promoter-driven *At*NRG1.1-nVenus and/or *At*NRG1.1-cCFP, *At*EDS1-FLAG, *At*SAG101-FLAG, or GUS-nVenus as negative control. Data collected in three independent experiments. One biological replicate is shown as representative.

**Fig. S3.**
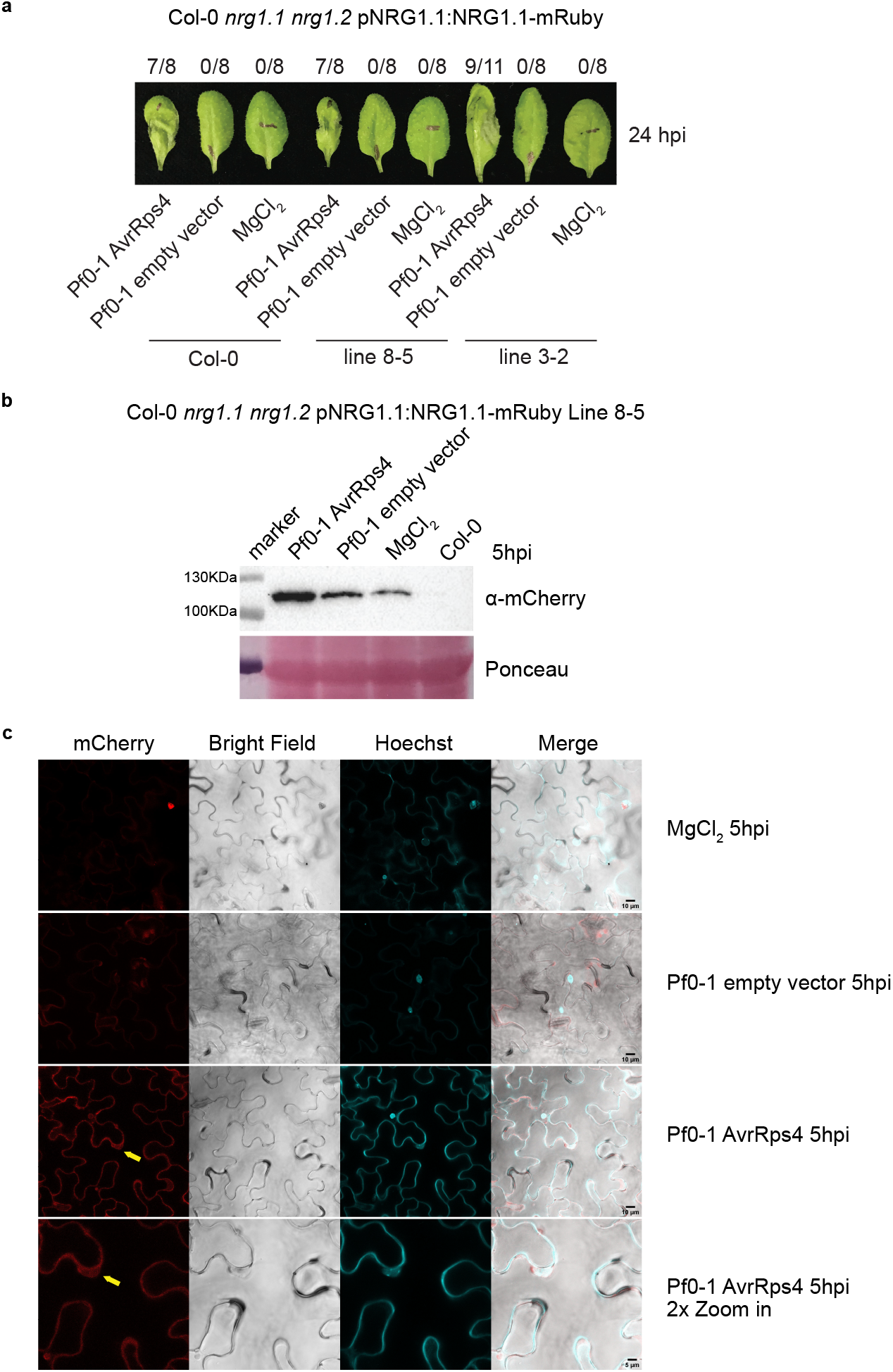
NRG1-mRuby is localized to nuclei upon AvrRps4 delivery by Pf0-1. **(a)** Macroscopic cell death in leaves of two independent lines of Col-0 *nrg1*.*1 nrg1*.*2* carrying pNRG1.1:NRG1.1-mRuby 24 hpi of Pf0-1 AvrRps4. Wild type was used as positive control. Infiltration of Pf0-1 EV or MgCl_2_ served as negative controls. Numbers in parentheses indicate leaves showing visual tissue collapse/total infiltrated leaves per genotype. Data was collected in three independent experiments. Representative images of one biological replicate are shown. **(b)** Higher accumulation of NRG1.1 is detected upon effector delivery in Arabidopsis. Leaves Col-0 *nrg1*.*1 nrg1*.*2* carrying pNRG1.1:NRG1.1-mRuby were infiltrated with Pf0-1 AvrRps4, Pf0-1 EV, or MgCl_2_. In three independent replicates, four leaf discs were harvested at 5 hpi for SDS-PAGE and Western blot analyses of NRG1.1 protein accumulation. **(c)** NRG1.1 localizes to PM and nuclei upon effector delivery in Arabidopsis. Live cell imaging of pNRG1.1:NRG1.1-mRuby stably expressed in Col-0 *nrg1*.*1 nrg1*.*2*. All images show single planes and micrographs were taken 4-5 hpi. Localizations were determined in three independent experiments and representative micrographs are shown. PM localization is observed in every cell and 2-4 nuclei could be imaged in one frame. Hoechst (cyan) and mCherry signal (red) are shown.

**Fig. S4.**
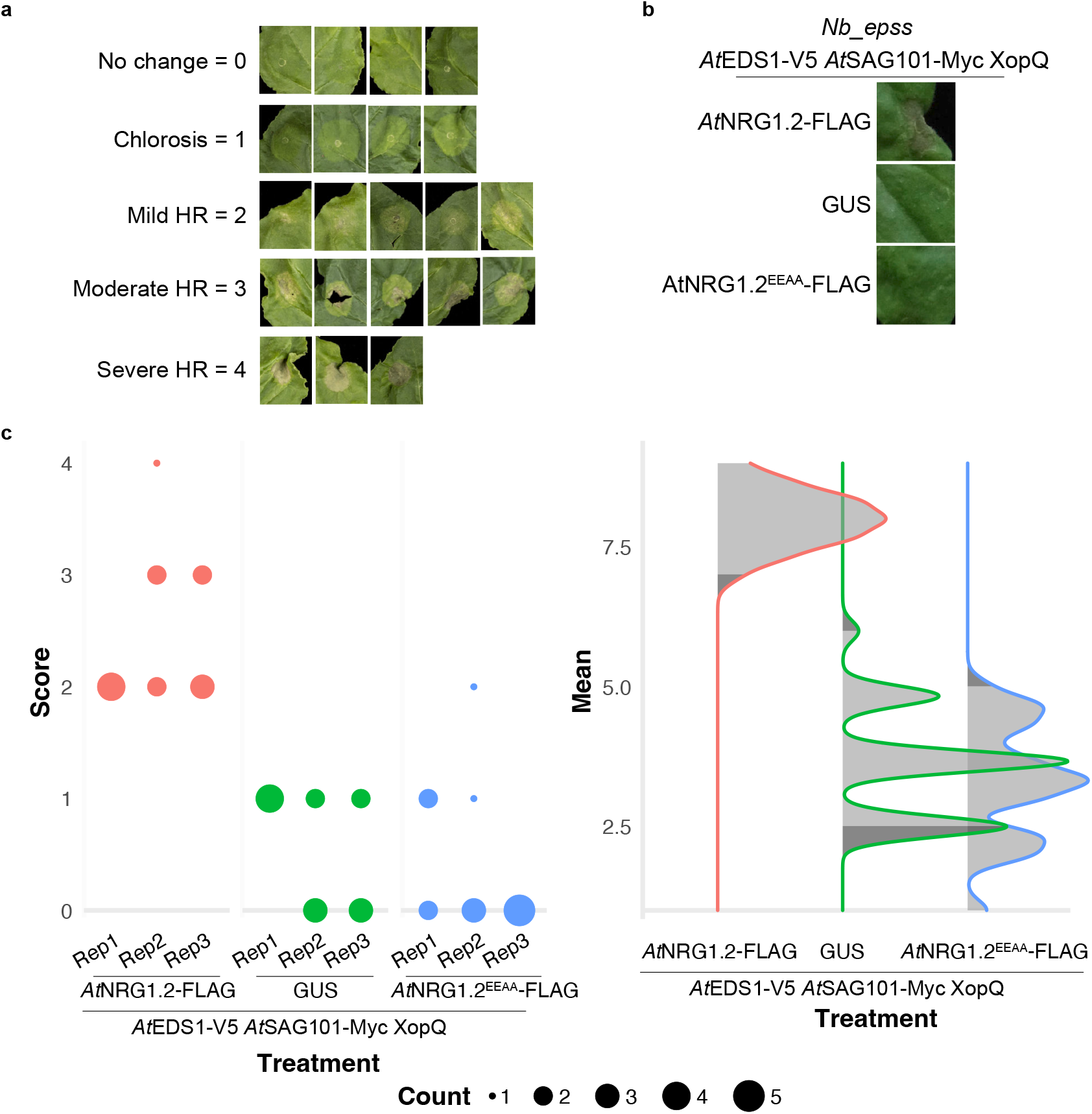
*At*NRG1.2^EEAA^-FLAG is defective in cell death assays in *N. benthamiana*. These data replicate what was observed for NRG1.1^EEAA^ in Sun., Lapin., Feehan et al. 2021 (12). **(a)** A bespoke cell death scoring system was generated for Agro-infiltration mediated reconstitution of Arabidopsis NRG1, EDS1, SAG101 in *Nb_epss* (11). A score of 0 (no change) and 1 (chlorosis) were not considered cell death. Representative images are shown from cell death assays in more than three representatives. **(b)** Representative images of one biological replicate from (c). Agro-mediated expression of 35S promoter-driven *At*EDS1-V5, *At*SAG101-Myc, and XopQ with *At*NRG1.2-FLAG, *At*NRG1^EEAA^-FLAG or GUS in *Nb_epss*. Cell death is visualized as tissue collapse and was scored 7 days post Agro-infiltration. **(c)** Replacement of glutamate residues with alanine in the N-termini of *At*NRG1.2^E21A,E29A^-FLAG results in loss of cell death phenotypes. Results are visualized in Derevnina plots (42). Dot plot size is proportional to sample number with confidence interval peaks adjacent. Experiment was performed with three independent biological replicates, each with 3-6 technical replicates. Estimation statistical tests were implemented with besthr R library (43). Bootstrapping resampling tests were performed with lower significance cut-off of 0.025 and upper of 0.975. Non-overlapping confidence interval peaks were considered significant.

**Fig. S5.**
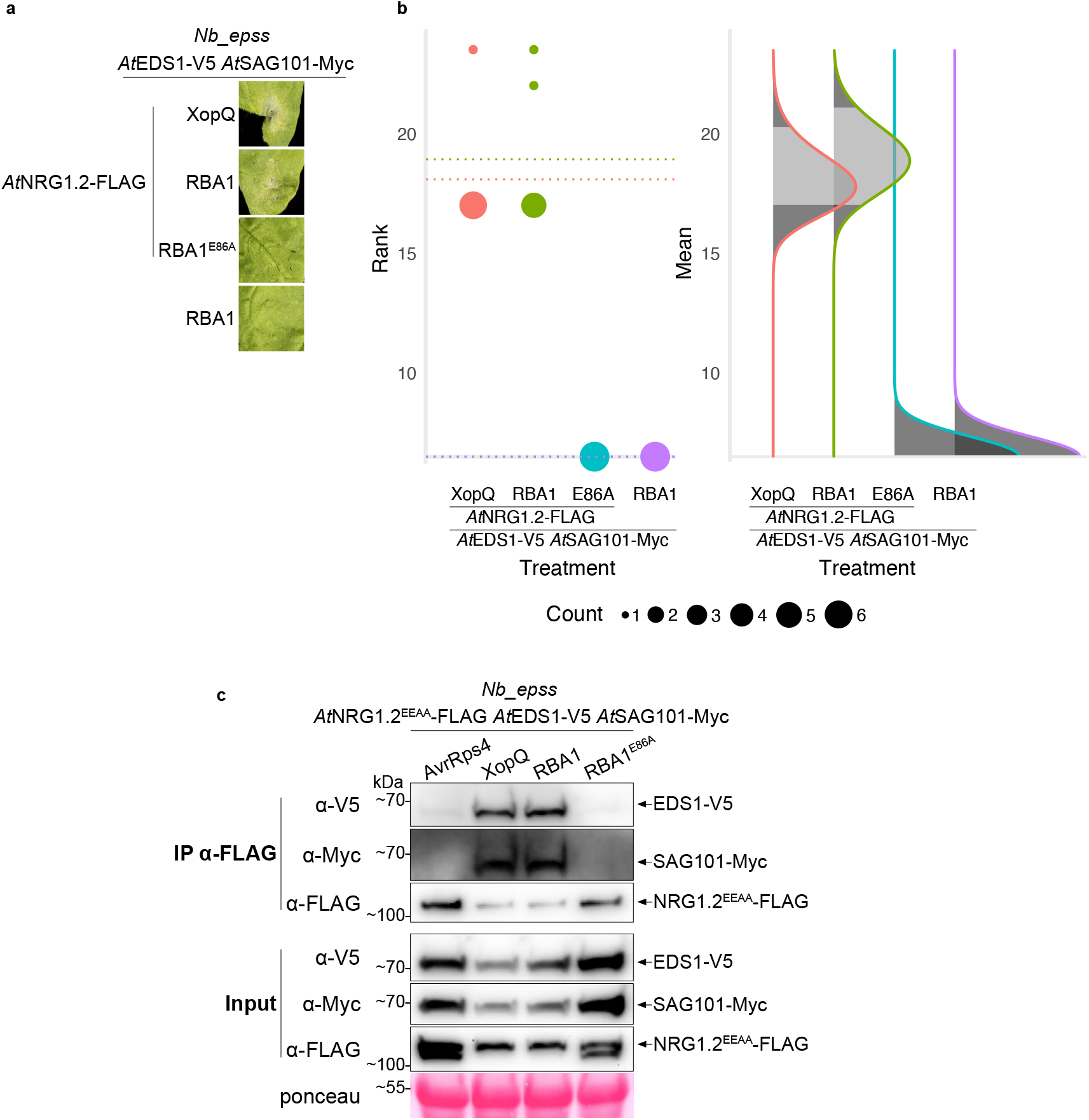
RBA1 induces EDS1 and SAG101 association with NRG1, which is lost when co-expressed with RBA1^E86A^. **(a)** Representative images of one biological replicate from (b). Agro-mediated expression of 35S promoter-driven *At*EDS1-V5 and *At*SAG101-Myc, with or without *At*NRG1.2-FLAG, and XopQ, RBA1, or RBA1^E86A^ in *Nb_epss*. Cell death is visualized as tissue collapse and was scored 7 days post Agro-infiltration. **(b)** Replacement of glutamate residue with alanine in RBA1^E86A^ does not induce cell death in Arabidopsis NRG1, EDS1, SAG101 reconstitution assays (11, 29). Results are visualized in Derevnina plots (42) with dot plot size proportional to sample number and with adjacent confidence interval peaks. Estimation statistical tests were implemented with besthr R library (43). Bootstrapping resampling tests were performed with lower significance cut-off of 0.025 and upper of 0.975. Non-overlapping confidence interval peaks were considered significant. **(c)** Replacement of glutamate residue with alanine in RBA1^E86A^ abolishes the RBA1-induced association of EDS1 and SAG101 with NRG1.2^EEAA^. SDS-PAGE and Western blot of coIP after Agro-infiltration and transient expression of 35S promoter-driven *At*NRG1^EEAA^-FLAG, *At*EDS1-V5, and *At*SAG101-Myc with AvrRps4, XopQ, RBA1, or RBA1^E86A^ in *Nb*_*epss*. NRG1.2^EEAA^ was used to remove confounding variable of cell death. Tissue was harvested 48 hpi. Co-delivery of XopQ was utilized to activation TNL signaling and induce *At*NRG1.2^EEAA^-FLAG association with AtEDS1-V5 and *At*SAG101-Myc. AvrRps4 was used as a negative control for TNL activation. Experiments were performed on three independent biological replicates with similar results.

**Fig. S6.**
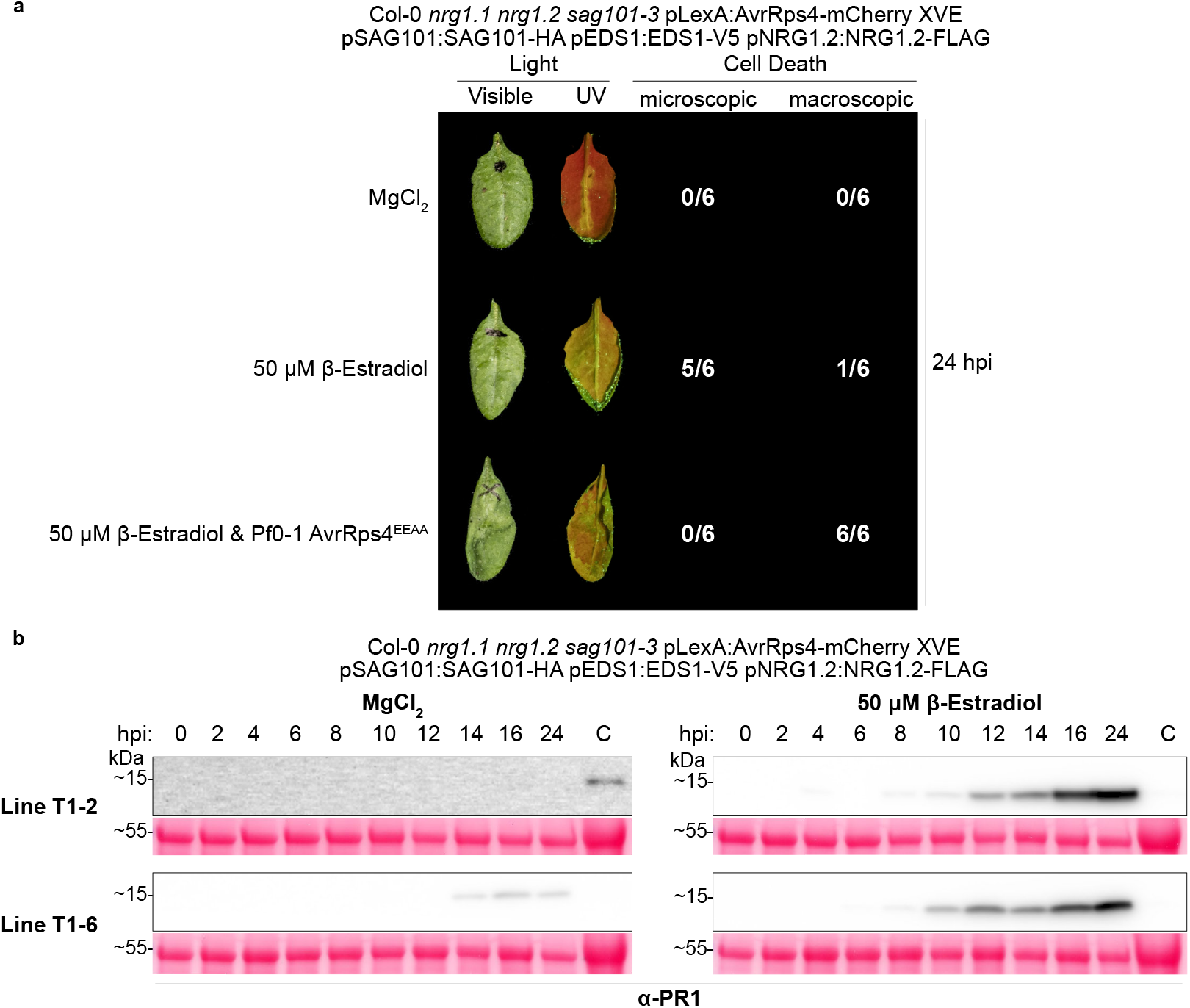
β-estradiol treatment in inducible lines reproduces microscopic cell death and defense gene activation. **(a)** β-estradiol-mediated induction of AvrRps4-mCherry with XVE system induces microscopic cell death, and β-estradiol co-infiltration with Pf0-1 AvrRps4^EEAA^ reconstitutes macroscopic cell death in stated Arabidopsis stable transgenic line (33). MgCl_2_ is used as a negative control. White light images were collected to demonstrate tissue collapse. Ultraviolet light images were collected to demonstrate auto-fluorescent cell leakage. Numbers in parentheses indicate number of leaves showing autofluorescence (microscopic) or visual tissue collapse (macroscopic)/total number of infiltrated leaves per genotype. Images were collected 24 hpi and a representative of one biological replicate is shown. Experiment was performed in two independent Arabidopsis stable lines to reconfirm Ngou et al. 2020 (33). **(b)** β-estradiol-mediated induction of AvrRps4-mCherry with XVE system induces defense activation (20). SDS-PAGE and Western blot of lysates for leaf protein accumulation assays. Defense activation was evaluated by immunolabelling for native α-PR-1 (Agrisera: AS10 687). MgCl_2_ was used as negative control. Leaves were painted to control for damage-induced PR-1 induction. “C” indicates positive control for α-PR-1 detection. Experiment was performed in two independent Arabidopsis stable lines to reconfirm Ngou et al. 2021 (20).

**Fig. S7.**
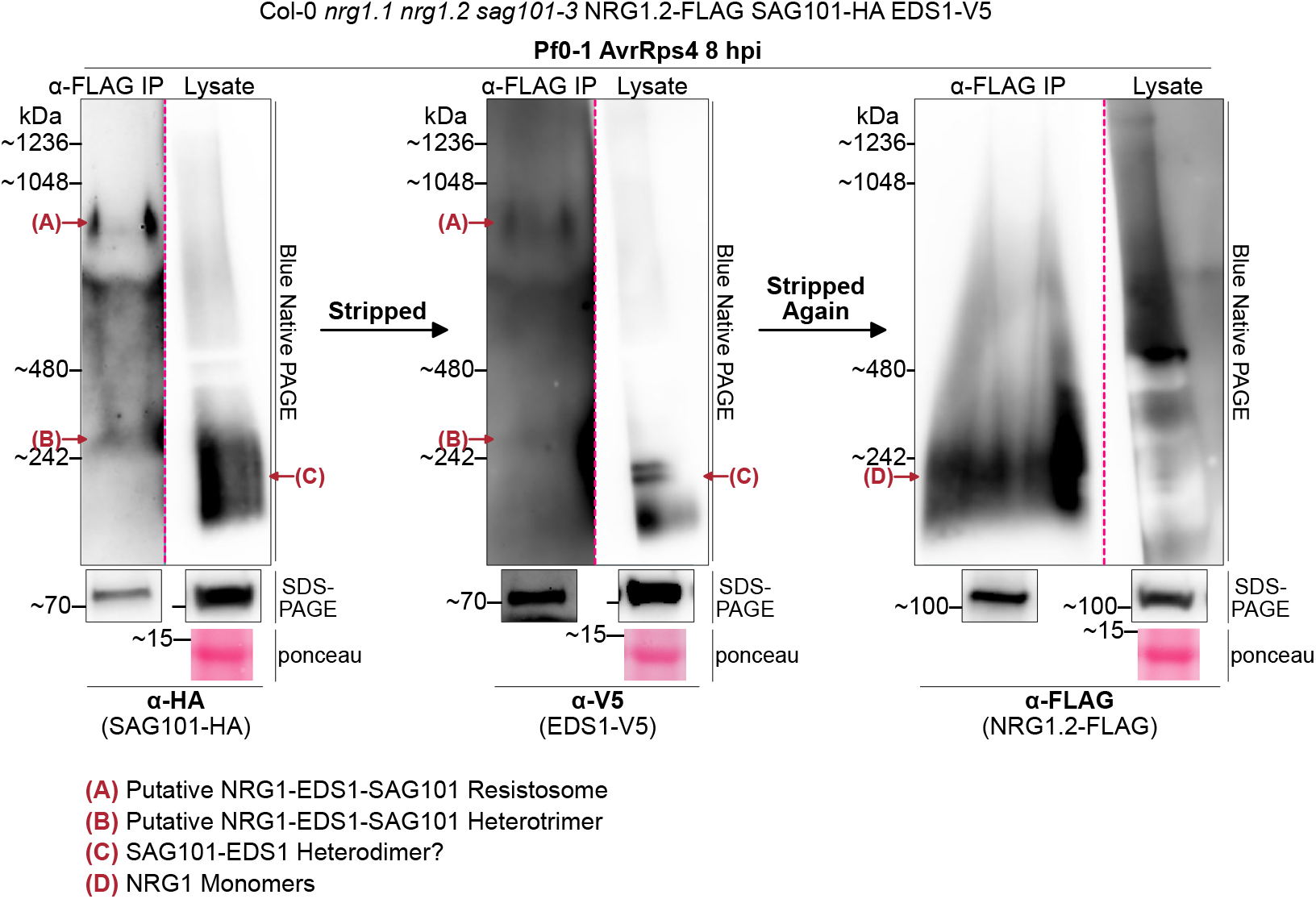
Faster migrating species of SAG101-HA and EDS1-V5 after α-FLAG IP of NRG1.2-FLAG in immune-activated tissues migrate slower than pre-activated EDS1-SAG101 heterodimers and NRG1.2 monomers. BN-PAGE and Western blot of lysates and coIP elution products from native promoter-driven stable Arabidopsis lines. The same membrane is shown after sequential immunolabelling and stripping with Restore^™^ Western Blot Stripping Buffer (21059). First immunolabelling was α-HA to detect SAG101-HA, second immunolabelling was α-V5 to detect EDS1-V5, and third immunolabelling was α-FLAG to detect NRG1.2-FLAG. Left lane shows elution product after α-FLAG IP of NRG1.2-FLAG while right lane shows lysate. Left lane is representative of post-activation complex formations for EDS1 and SAG101, and right lane is representative of pre-activation states for NRG1, EDS1, and SAG101 which do not change in lysates from mock/un-infiltrated to AvrRps4-treated (Fig. 3). Lysate in α-FLAG blot shows bleaching effects, likely an artifact of sequential stripping. In α-FLAG blot, α-FLAG IP shows pre-activation state of NRG1.2-FLAG as large NRG1 oligomer is not observed, and banding patterns do not change, from mock/un-infiltrated to AvrRps4-treated and observation of oligomeric NRG1 requires detection of NRG1.2-V5 after α-FLAG IP of NRG1.2-FLAG (Fig. 3a & 3c). Species identities are indicated in red, and red dashed lines indicate lanes were cropped. Samples were run in parallel on same membrane. Two independent biological replicates showed similar results.

